# Long-term shift in community composition of deadwood fungi after clear-cutting

**DOI:** 10.64898/2026.01.20.700536

**Authors:** Eivind Kverme Ronold, Håvard Kauserud, Jenni Nordén, Johan Asplund, Rune Halvorsen, Line Nybakken, Anders K. Krabberød, Inger Skrede, Sundy Maurice

## Abstract

1. Rising anthropogenic pressures in the 20th century have caused extensive habitat loss and fragmentation, threatening biodiversity across ecosystems worldwide. In the boreal forest of Fennoscandia, clear-cut forestry has been a major driver of these changes, resulting in a fragmented landscape of even-aged forest stands. Several studies in recent years have investigated the effect of habitat fragmentation and loss on fungal communities associated with deadwood, but these have mainly focused on visible sporocarps and mushrooms. The effects of forestry on the whole fungal community within deadwood have not been explored as extensively, and especially not in the context of long-term effects.
2. We investigated the effects of clear-cutting on deadwood-inhabiting fungi in boreal *Picea abies* forests in Norway using ITS2 metabarcoding of sawdust samples collected from 459 logs distributed across 24 paired near-natural and previously clear-cut plots harvested 50 to 90 years ago.
3. While the overall fungal richness associated with deadwood was similar between the two management types, community composition differed markedly. Plot-scale fungal diversity was linked to deadwood heterogeneity, and composition of the common species was additionally affected by the living tree structure. Nearly all red-listed species were exclusively found in the near-natural forest plots. These findings demonstrate that clear-cut forestry shifts fungal community structure rather than reducing the total number of species and that rare and common species are structured by different environmental drivers but respond in similar ways to large scale disturbance.
4. *Synthesis:* Our results highlight the importance of maintaining structural complexity of deadwood in boreal forests for fungal conservation. We found a consistent relationship between deadwood volumes and heterogeneity, and community diversity and species richness. This relationship was consistent even for previously clear-cut forests, showing that retaining structural heterogeneity in managed ecosystems has a positive impact on species diversity.

## Introduction

Increasing pressure from anthropogenic activities through the 20^th^ century has caused widespread habitat loss and fragmentation across the globe, and is a major threat to biodiversity (e.g., Johnson et al., 2017; Díaz et al., 2019; Gonçalves-Souza et al., 2025; Keck et al., 2025). Fennoscandia has a long history of forestry, and since the end of the Second World War, stand-based forest management with clear-cut harvesting followed by tree planting has been the pervasive method of management (Kuuluvainen et al., 2012). Consequently, the forest landscape is fragmented, with a mosaic of even-aged stands in different successional stages. In southeastern Norway, 53-58% of the productive forest, and 85-90% of the high-yield spruce forests, have gone through one cycle of clear-cutting (Viken, 2021). Most of the remaining forests were subjected to selective logging prior to 1940. Consequently, less than two percent of the forests can be considered pristine, with negligible history of anthropogenic influence (Storaunet and Rolstad, 2020). Numerous studies have documented the negative effects of clear-cut forestry on richness and diversity of various organismal groups (Lunde et al., 2025; Siitonen, 2001; Work and Hibbert, 2011), including fungi inhabiting deadwood (Junninen and Komonen, 2011; Lunde et al., 2025). The majority of terrestrial species under threat and on the Norwegian red-list are associated with old-growth forests (Artsdatabanken, 2021).

Despite a large body of evidence highlighting the importance of biodiversity for ecosystem functioning and services (Cardinale et al., 2012), defining and quantifying diversity remains challenging (Fletcher et al., 2025; Mori et al., 2018; Socolar et al., 2016). Traditional measures of species richness (alpha-diversity) have been useful for elucidating the relationships between the number of species and community response to disturbances (Lunde et al., 2025; Lundström et al., 2013; Stenbacka et al., 2010; Viljur et al., 2022). However, it is becoming increasingly clear that species richness alone is insufficient to capture key changes in biodiversity under natural or anthropogenic disturbances, as it is disconnected from species turnover (Bratli et al., 2006; Fletcher et al., 2025; Nirhamo et al., 2025), which is a highly relevant consideration in conservation biology (Gonçalves-Souza et al., 2025; Socolar et al., 2016).

The compositional difference between sites (beta-diversity) is a powerful tool for describing community responses to disturbances (Rydgren et al., 2019; Socolar et al., 2016). Partitioning beta-diversity into components of species turnover between sites and the degree of nestedness, i.e., whether one site mostly contains a subset of the community in a more species-rich site (Baselga, 2010), can be especially useful when comparing a disturbed habitat with a less disturbed reference habitat. As an extension to beta-diversity which strictly compares communities in a pairwise fashion, the concept of zeta-diversity has been introduced (Hui and McGeoch, 2014; Latombe et al., 2017). By extending the concept of pairwise community dissimilarity into multi-site comparisons, this approach to biodiversity enables modelling the effects of disturbance separately on rare and common species (e.g., Erős et al., 2020; Parra□Sanchez et al., 2025). This is useful in conservation ecology, as habitat fragmentation and disturbance has been shown to differently affect rare and common species (e.g., Erős et al., 2020; Nordén et al., 2013; van Galen et al., 2023).

The ecological effects of clear-cutting as a harvest method deviate significantly from those of natural stand-replacing disturbances such as forest fires and windthrow, in terms of removal of biomass, soil compaction, and regeneration by planting of one tree species that later forms the dominant canopy (Kuuluvainen, 2009; Siitonen, 2001). Consequently, much of the naturally structured and dynamic habitats with deadwood in various stages of decay are reduced or removed entirely (Asplund et al., 2024; Kuuluvainen, 2009; Kuuluvainen et al., 2012). The amount of deadwood on the forest floor is significantly reduced in managed forests (Asplund et al., 2024; Jonsson et al., 2016; Lunde et al., 2025; Siitonen, 2001). This reduction of deadwood volumes and the loss of large deadwood units in particular may have strong negative impacts on organisms that depend on the continuous presence and turnover of deadwood in the forest (Stokland et al., 2012).

Several organismal groups are dependent on deadwood as the primary habitat, including e.g., wood-decaying fungi (Leonhardt et al., 2019; Rayner and Boddy, 1988; Rustøen et al., 2023). Additionally, many organismal groups use deadwood during part of their life cycle (Stokland et al., 2012), including e.g., many insects (Hammond et al., 2004; Work and Hibbert, 2011) and amphibians (Margenau et al., 2023). Wood-decaying fungi are especially important organisms associated with deadwood, since they are the main drivers of wood decomposition (Rayner and Boddy, 1988). Fungal communities inhabiting deadwood display strong successional patterns throughout the decay process (Mäkipää et al., 2017; Rajala et al., 2012, 2015). Initial colonization by pioneering Basidiomycota species, together with Ascomycota soft-rot species initiates the decomposition (Rajala et al., 2015; Rayner and Boddy, 1988; Schilling et al., 2020), potentially along with species living as endophytes within the bark and wood before the tree dies (Griffith and Boddy, 1990; Saine et al., 2024; Song et al., 2017). Physical and chemical changes to the wood, along with species interactions, drive community succession resulting in turnover of species adapted for deadwood with certain characteristics (Nenzén et al., 2025; Ottosson et al., 2014; Rajala et al., 2015b). Fungal diversity in the decomposing wood tends to increase over time, as many early colonizers stay active while new species continue to establish (Mäkipää et al., 2017). Species in the Basidiomycota genera *Phellinus* and *Antrodia* are typically abundant in intermediate stages of coniferous wood decay (Rajala et al., 2015). The red-listed species *Phellopilus nigrolimitatus* and *Fomitopsis rosea* are often associated with intermediate- to late-decay deadwood in Fennoscandian forests (Stokland and Kauserud, 2004; Artsdatabanken, 2021). As the decomposing wood become more integrated in the soil, many litter saprotroph and ectomycorrhizal species are often found as part of the community inside the wood substrate, reflecting a more gradual transition between soil and deadwood fungal communities (Mäkipää et al., 2017). Depending on the size of the deadwood, regional and local site conditions and climatic variables, complete decomposition of coniferous deadwood in Fennoscandian boreal forests may take a century or more (Niemelä et al., 2002; Tuomi et al., 2011).

The characteristics of the deadwood play a critical role for the biodiversity associated with the substrate. Different tree species, sizes, and diameters attract different organisms and host different fungal communities (Purahong et al., 2016; Stokland et al., 2012; van der Wal et al., 2015). Many of the red-listed fungi in Fennoscandian forests are associated with large units of deadwood and depend on the continued presence of naturally decaying wood or old large living trees (Artsdatabanken, 2021; Stokland and Kauserud, 2004; Tikkanen et al., 2006). Harvesting may cause a short-term spike in species richness, as harvest residue may serve as an abundant and diverse habitat for a decade or more (Suominen et al., 2019). However, there is a lack of long-term studies on diversity beyond the first decade (highlighted in Lunde et al., 2025). Moreover, studies of deadwood-decaying fungi have mainly focused on fungi producing macroscopic sporocarps (Juutilainen et al., 2014; Moor et al., 2021; Runnel et al., 2021; Suominen et al., 2019). However, other inconspicuous fungi are also important players during the decomposition process (Parfitt et al., 2010), including yeasts and many species in Ascomycota. These saproxylic fungi have been less studied in the context of conservation ecology, but the richness and diversity in these communities have been shown to be higher in natural forests (Purhonen et al., 2021). A recent DNA-based study in Finland found that naturally fallen deadwood hosted richer and more diverse communities of pioneering fungi (in the orders Polyporales, Hymenochaetales and Helotiales) than artificially restored (i.e., felled and placed into the forest) deadwood (Saine et al., 2024). By describing a much larger part of the community, environmental DNA methods are useful tools in conservation ecology to investigate the effects of disturbance on fungal communities (Heilmann-Clausen et al., 2015).

In the current study, we aim to assess the long-term effects of clear-cutting on deadwood-inhabiting fungal communities in boreal forests in southeastern Norway. This aim was addressed by collecting samples from deadwood in paired *Picea abies* forest plots, where each of the 12 sites consisted of a mature, previously clear-cut plot and a near-natural plot with no history of intensive management. We hypothesize (H1) that the near-natural plots will have a higher fungal species richness and diversity than the previously clear-cut plots due to larger volumes and diverse quality of deadwood. Additionally, we expect (H2) that the near-natural plots, due to higher volume and variability of deadwood, will host more red-listed species than the previous clear-cut plots. We also expect (H3) species turnover (i.e., species replacement) to be the main driver of differences between forest pairs, not nestedness (i.e., species loss/gain). Lastly, we posit (H4) that rare species (low zeta-order) will be more strongly impacted by the change in dead wood diversity than intermediate and common species (higher zeta-order).

## Materials and methods

### Study plots

In this study we used a set of 12 paired Norway spruce (*Picea abies*) dominated forest sites (Fig. 1a). Each site-pair consisted of one plot in a previously clear-cut forest stand (hereafter CC) (stand age ranging from 50 - 90 years at time of sampling in 2022) and one plot in a nearby stand that had remained in a near-natural state (hereafter NN) exhibiting minimal human impact. We ensured that the forest plots within each pair were as similar as possible in physical and topographic characteristics, including elevation, slope, aspect, edaphic factors, and site index (Supplementary methods). The average distance between each pair of CC and NN was 1270 m (range 540 – 3140 m). Within each of the 24 plots, we established a 15 × 133.3 m (0.2 ha) transect (Fig. 1b) for surveying deadwood. In addition, within each transect, a 15 × 15 m central study area was established in which we registered representative structural properties of the forest plot; the density and species of living trees, the volume of living spruce (calculated following Vestjordet, 1967), and the coefficient of variation in diameter at breast height (DBH) for all standing spruce (Fig. 1c). A detailed description of the 24 plots and the study design can be found in Asplund et al. (2024).

**Figure 1:**
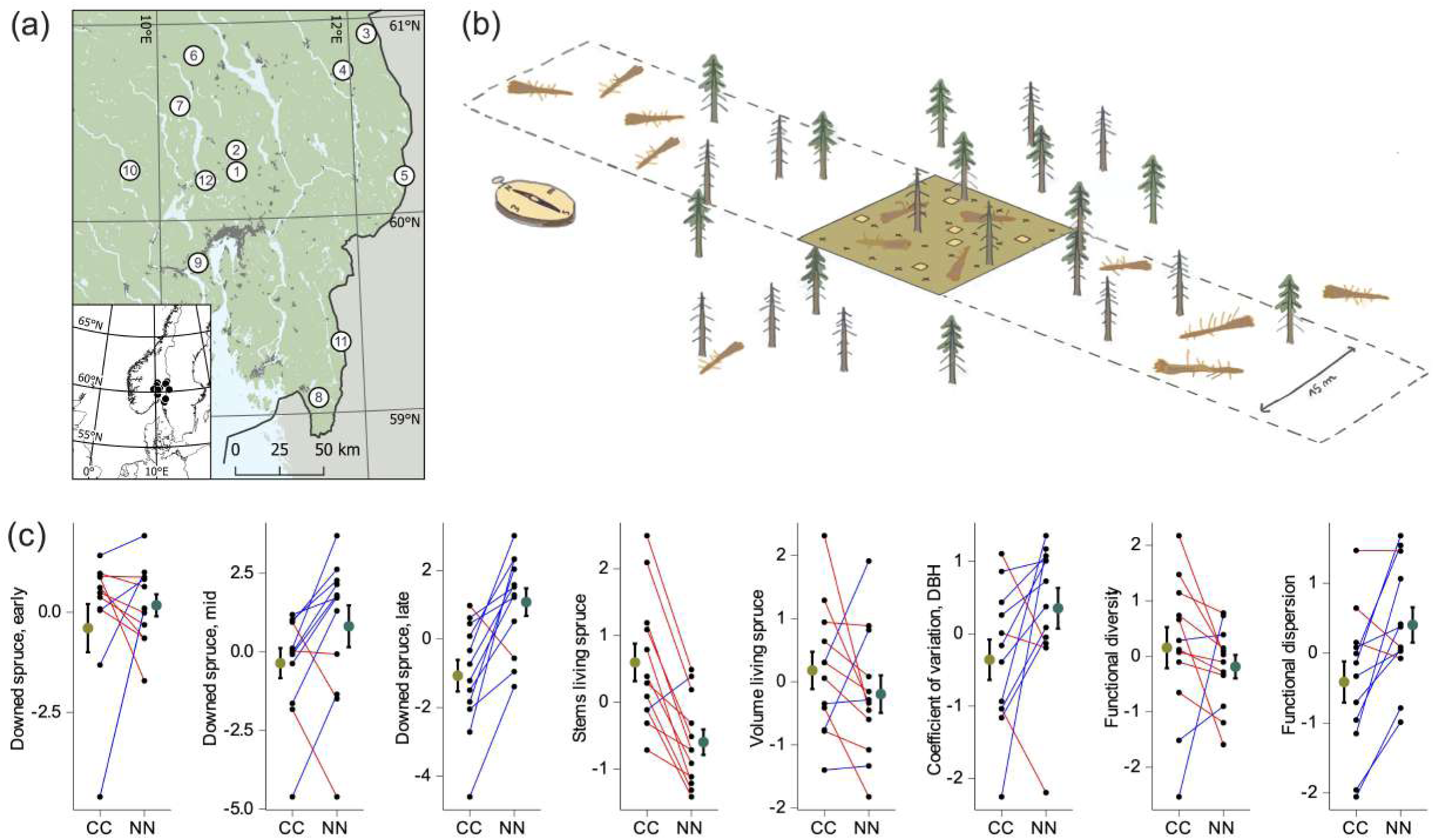
(a) Map of the twelve study sites in south-eastern Norway, one near-natural (NN) and one previously clear-cut (CC) site was established at each location. (b) Illustration of the transect design in the study sites. Each transect is 133 m × 15 m, with a central 15 m × 15 m plot. We surveyed all deadwood logs with a diameter ≥5 cm that had their origin within the transect and collected sawdust samples for eDNA analysis from a randomized subset of 20 spruce logs. The central plot was used for assessing forest structural characteristics such as stem density, living tree volume and variation in DBH of the living trees. (c) Overview of a selection of forest and deadwood properties and the difference between NN and CC plots, scaled to zero mean and unit variance. Lines connect the paired plots; blue lines indicate higher value in the NN plot and red lines indicate higher value in the CC plot.

### Deadwood survey and sampling

We surveyed all downed deadwood (logs) with DBH or basal diameter (when the breast height point was not present) ≥ 5 cm, provided the point of origin was within the transect. For detailed descriptions of the deadwood survey we refer to Asplund et al. (2024). We subsequently selected a random subset of 20 spruce logs (18 from Hemberget with less deadwood) within each transect by attributing 20 random numbers from the list of all surveyed dead wood for each plot. One plot (Hemberget CC) was surveyed in the entire stand, not just the transect due to very low amounts of deadwood present.. The percentage of logs present in each plot sampled ranged from 17.5% to 100% (Table S1). From each of these 20 logs, we collected sawdust into Ziplock bags using a Makita cordless drill (model DDF484RTJ) and a 22 mm drill bit. We first shaved off the bark and the outer 2–5 mm of wood by use of a knife. We drilled holes perpendicular to the ground at approximately 6 cm depth, or to the opposite end without penetrating the log for those with smaller diameter. To ensure that we collected a representative amount of sawdust from each log, we drilled one hole every 2.5 m along the log starting at 0.5 m distance from the base of the log (or 0.5 m in from the root collar uprooted logs) to avoid the break point or the root collar. For logs shorter than 2.5 we drilled the sample from the centre along the length of the log. All logs were drilled from only one side and the sawdust collected per log was pooled in one Ziplock bag. To reduce contamination, the knife and the drill bit were brushed with a steel brush and flamed using 80% ethanol after sampling each log. The sawdust samples were stored in a cooler with icepacks while working in the field for a day or up to 3 days, then placed at –20 °C awaiting further processing. In total, we sampled 471 unique dead wood units across the 24 studied plots. Selecting only 20 logs, regardless of total number of logs per site, differ from many other sporocarp based surveys where all the logs in an area are investigated for target species groups. Our randomized selection ensured that we sampled a representative subset of the deadwood present per plot (Fig S1).

### Sample processing

The deadwood samples were homogenized by thoroughly mixing the sawdust within each bag. A 50 ml Falcon Tube with known weight was filled with approximately 6 g of fresh sawdust (all weights were recorded with precision ± 0.01 g). The tubes were covered with absorbent paper and then freeze-dried (GAMMA 2-16 LSC) for 48 h at 0.5 mbar. Tests showed that no further weight loss took place after 48 h.

The moisture content of the sawdust was measured by weighing the samples before and after freeze-drying. The dried sawdust was then evenly split into two 50 ml Falcon Tubes: one for the biochemical analyses and the other for the DNA analyses, respectively. The sawdust intended for DNA isolation was crushed by adding two 5 mm diameter ceramic beads into the tube and crushing the tubes in a FastPrep 24 (MP Biomedicals, USA) at a speed of 4.5 m/s for 45 s. The crushing process was repeated at least three times, or until the sawdust was fully pulverized.

### DNA extraction, library preparation, and sequencing

We extracted DNA from the second aliquot of dried and crushed sawdust using an in-house CTAB/chloroform protocol with an additional DNA-column cleaning step.

DNA extracts were cleaned using an E.Z.N.A 96-well plate kit following the manufacturer’s protocol (Omega Bio-tek). The filtrate collected in 96-well PCR microplates was stored at –20°C.

We generated fungal ITS2 libraries using primer fITS7 (GTGARTCATCGAATCTTTG) (Ihrmark et al., 2012) and ITS4 (TCCTCSSCTTATTGATATGC) (White et al., 1990), with the reverse primer modified by two degenerate bases (S at positions 6 and 7) to reduce mismatches to certain Ascomycota species e.g., Archaeorhizomycetes. Each library was multiplexed with 96 uniquely tagged primer pairs. PCR reaction was performed in 25 μl volume consisting of 15.75 μl distilled water, 5 μl 5x concentrated Q5 buffer, 1.25 μl of each primer at 10 mM, 0.25 μl Q5 DNA polymerase, and 1 μl DNA template. Cycling conditions were 95°C for 5 min, followed by 32 cycles of 95°C for 45 s, 56°C for 45 s, 72°C for 60 s and a final extension step at 72°C for 10 min. We evaluated the PCR amplifications from the six PCR plates on a 1% agarose gel.

The DNA concentration in each well was normalized using the SequelPrep Normalization Plate kit (Thermo Fischer Scientific) following the manufacturer’s protocol. After normalization, each 96-well plate was split into three pools for AMPure bead purification (Beckman Coulter Inc., Brea, CA, USA). Within each of these new pools, we performed an AMPure XP beads purification to remove primer-dimers and other smaller DNA fragments following an in-house protocol. We eluted the DNA in 20 μl of elution buffer and finally pooled it into 60 μl of ITS2 amplicons for each of the PCR libraries and measured the amplicon concentrations using Qubit fluorescence spectrophotometer (Life Technologies). The six ITS2 libraries were barcoded with Illumina adapters and paired-end sequenced on two Illumina MiSeq lanes (with V3 chemistry) at Fasteris (Plan-Les-Ouates (GE), Switzerland).

### Bioinformatics analyses

Bioinformatics analyses were performed on the SAGA computer cluster provided by Sigma2 – the National Infrastructure for High-Performance Computing and Data Storage in Norway, using a custom pipeline tailored for ITS2 amplicon sequences. Paired-end reads were demultiplexed and primers removed using Cutadapt v4.2 (Martin, 2011). The demultiplexed samples were denoised, chimera checked, and clustered using DADA2 (Callahan et al., 2016) with global option *BAND_SIZE=32* to account for length variation in the ITS2 sequences. The *filterAndTrim* function *maxEE=c(2,2)* allowed 2 errors per direction when aligning sense and antisense sequences, and *trimRight=10* to remove 10 nucleotides at the end of each sequence as these are generally of lower quality. The *learnErrors()* function was trained on 5×10^8^ nucleotides after checking that the error models from the default 1×10^7^ were not producing satisfying error plots.

OTUs generated from DADA2 were trimmed using ITSx (Bengtsson□Palme et al., 2013) and the resulting OTUs were further clustered using VSEARCH v.2.22 (Rognes et al., 2016) at 97% similarity. A second *denovo* chimera check was run with VSEARCH using options *--uchime-denovo* and *--nonchimeras* to ensure removal of any residual chimeras. The OTU table was further curated using the mumu v1.0.2 post-clustering algorithm (https://github.com/frederic-mahe/mumu) to account for over splitting of OTUs and the high intraspecies variation in fungal ITS2. Taxonomic annotation was performed with SINTAX (Edgar, 2016) as implemented in USEARCH v11.0 (Edgar, 2010) against a trimmed version of the UNITE database (Nilsson et al., 2019, version 16.10.2022) in which all taxa headers containing either “Incertae_sedis” at phylum and class rank or an “unknown” classification at any rank were removed before assignment. The resulting OTU table consisted of 4,238 OTUs across 567 sawdust samples, including controls and replicates. The pipeline and the scripts are available on GitHub: https://github.com/ekronold/Zazzy_metabarcoding_pipeline.

### OTU table curation

The OTU table was further curated in R v4.5.0 (R Core Team, 2024). First, all the positive and negative controls were investigated and 51 OTUs in the controls were removed. Due to the large number of samples spread across six 96-well plates, we included replicate samples both within each plate and across plates, totalling 121 replicate samples. These were checked for similarity using DCA ordination with the function *decorana()* from the package *vegan* (Oksanen et al. 2025). The replicate sample with the highest number of reads was retained in the OTU table. The curated OTU table consisted of 470 samples with mean sequence reads per sample of 24,042 (range 76–265,140) and 156 mean OTUs per sample (range 21–427). We further rarefied the OTU table to 1000 sequence reads using the function *rrarefy()* in *vegan* (Oksanen et al. 2025), discarding 11 low-read samples from further analyses. To compare the fungal communities between plots, we transformed the rarefied OTU table into a presence/absence matrix and summed occurrences per OTU per plot, giving plot-level observation counts (of max. 20 for all sites but one with a max. of 18) for each OTU. This final OTU table with 24 plots and a 0-20 (18) count value per OTU was used for all downstream analyses.

### Predictor variables

To use deadwood diversity as predictors in our models, standard functional diversity indices were calculated using the package mFD (Magneville et al., 2022) by treating the categorical variables ‘tree species’, ‘decay stage’ (on a 1–5 scale) and type (e.g. uprooted, man-made or broken) and the continuous variables percentage of ‘bark cover’, ‘epiphyte cover’, ‘ground contact’ and ‘volume’ in m^3^ as traits (Fig. 1C). A full description of the deadwood metric calculation is presented in supplementary methods (Fig. S1) and in Asplund et al.,(2025). In short, four indices were calculated: functional richness (FRic), functional evenness (FEve), functional dispersion (FDis) and functional diversity (FDiv). Each metric was calculated from a multivariate ordination of all the deadwood units based on the above deadwood properties. The four metrics describe different aspects of the available habitat represented by the variety of deadwood in the plots. FRic represents the extent of the multivariate space and will mostly be driven by the number of deadwood logs. The values of FEve will be higher if the deadwood in a plot consists of an even distribution of logs across the functional trait space, indicating a more regular representation of different niches. FDiv represents how much the functional composition of logs in a plot differs from the average composition of log properties in the plot. Lastly, FDis represents how much each single log in the ordination differs from the average log in the plot. The latter two (FDiv and FDis) are two different ways of representing substrate heterogeneity in the plot.

The number, species and DBH of all living trees in the plot were registered. Standard bioclimatic variables (Fick and Hijmans, 2017) were derived from monthly temperature and precipitation data from seNorge (v.2) on a 1-km grid (Lussana et al., 2019). All data were adapted to a 100 × 100 m grid using ordinary kriging in SAGA GIS v.2.3 (Conrad et al., 2015) or rasterization. We also calculated descriptive community indices (richness, Shannon-diversity, and beta-dispersion) for each transect using the package *vegan* (Oksanen et al., 2025). Richness was calculated simply as the number of OTUs using the function *specnumber()*, Shannon-diversity was calculated using the function *diversity()* with the option *index = “Shannon”*. Beta-dispersion was calculated by first generating a Bray-Curtis distance matrix of the rarefied OTU table containing all 460 deadwood logs with the function *vegdist()*. We then applied the function *betadisper()* on the distance matrix using the plot ID as a grouping variable. The beta-dispersion values were calculated as the median distance to the group centroid of each sample within each group.

### Statistical analyses

All statistical analyses were performed in R v4.5.0 (R Core Team, 2024). To test for compositional variation related to the forest management (CC / NN), we first fit a PERMANOVA analysis by applying the *adonis2* function from vegan (Oksanen et al. 2025), restricting the permutations to between logs from paired plots from the same site. We generated a taxonomic tree using the package *metacoder* (Foster et al., 2017), with taxa truncated at family rank for better visual presentation. The function *compare_groups()* was used to calculate differences in OTU occurrences between the two forest types (CC and NN).

We calculated pairwise dissimilarity between plots with the function *decostand()* from package vegan (Okasenen et al. 2025) applying the Bray-Curtis distance to the summed OTU table. The community dissimilarity was analysed by fitting an NMDS ordination using the function *metaMDS()* with two dimensions and a minimum 1000 starting configurations from *vegan* (Oksanen et al. 2025). We then fit the environmental variables to the resulting ordination using the function envfit() from *vegan*. We summarized the fungal species richness within NN and CC plots for a selection of taxonomic and functional fungal OTU groups, as classified in FungalTraits (Põlme et al., 2020): “Saprotrophic, filamentous, Basidiomycetes”, “Saprotrophic, filamentous, Ascomycetes”, “Ectomycorrhizal”, “Yeasts”, “Dimorphic Yeasts”, and “Lichenized”. We also matched the species hypothesis from the taxonomic annotation to the list of red-listed wood-associated species in Norway (Artsdatabanken, 2021). We did this by checking the “Species hypothesis” annotated by SINTAX only for annotations with a bootstrap score of ≥0.8. Pairwise beta diversity separated into components of nestedness and turnover was calculated using the *beta.pair.abund()* function in the package *betapart* (Baselga et al., 2023). The beta-partitioning was visualized with a stacked bargraph generated using the *ggplot2* package (Wickham et al., 2019).

### Diversity models

To analyze the main drivers of community differences, we applied a generalized dissimilarity model using the package *gdm* (Mokany et al., 2022). The plot-pair object for the gdm model was prepared using the function *formatsitepair()* with Bray-Curtis distances. The pairwise distances were weighed by the number of OTUs per site, setting the option *weightType = “richness”,* as recommended for studies of presence-only data. We performed forward model selection among the environmental predictors, always including geographic distance by providing the latitude and longitude of the plots in the *formatsitepair()* function and fitting the option *geo=TRUE* in all models. Significant variables were selected based on p-values estimated using the function *gdm.varImp()*.

To further investigate the influential parameters on the rare and common parts of the species communities, we applied multisite generalized dissimilarity models using the function *zeta.msgdm()* in the package *zetadiv* (Latombe et al., 2017). We fitted models for the zeta-orders 2 through 10. We considered zeta-orders 2 and 3 as the rare community members, orders 4 through 6 as the intermediate community members and the remaining orders 7 to 10 as the common community members. For each predictor the model estimated I-splines using a Tweedie distribution with a log link by setting *reg.type=”ispline”*. For each rarity class, we incrementally increased the Tweedie p-parameter to optimize the model fit. Model fit was evaluated using the *appraise()* function in the package *gratia* (Simpson, 2024) to control for approximate normality. As the number of shared OTUs for each zeta-order depended on the number of random samples drawn from the data, we ran each model 50 times from different starting points, each time making 1000 random samples (Parra□Sanchez et al., 2025). For each zeta-order, we fitted a model using the full table of environmental predictors and checked for significance using the *summary()* function, using a median p-value of 0.05 across the 50 samples as the significance threshold. Non-significant predictors were removed, and the final model was fitted using only significant predictors, before extracting the I-splines using the *Return.ispline()* function. We extracted the I-spline from all 50 models and calculated the median values for visualization.

## Results

### OTU diversity and distribution

The rarefied OTU table consisted of 2,909 OTUs across sawdust samples from 459 logs, including 232 samples from near-natural (NN) forests and 227 from previously clear-cut (CC) forests. On average, 61 OTUs (range 8–129) were detected per sample; with a mean of 62 OTUs (range 14–129) in the NN plots, and 61 OTUs (range 6–123) in CC samples. The species accumulation curves did not saturate at 20 samples for any plot, however, the species accumulation curves for the entire CC and NN sample set showed indications of saturating at the total of 240 samples (20 logs in each of the 12 plots; Fig S2). Several fungal orders differed in their proportional distribution between the NN and CC plots (Fig. 2, Fig. S3). Among the OTU-rich orders, Polyporales and Hymenochaetales in Basidiomycota, as well as Helotiales, Umbilicariales and Sordariales in Ascomycota were most abundant in NN plots. In contrast, Russulales, Agaricales and Thelephorales in Basidiomycota, and Hypocreales, Saccharomycetales, Xylariales and Dothideales in Ascomycota were more abundant in the CC plots (Fig. 2). At finer taxonomic resolution, however, several families within the highly diverse orders e.g., Polyporales, Agaricales, Helotiales showed opposite preferences for NN vs. CC plots (Fig. 2). We also identified a high abundance of yeasts from the order Saccharomycetales. Among the ten families observed from Saccharomycetales, five occurred proportionally more in the CC plots and one was proportionally more abundant in NN.

**Figure 2.**
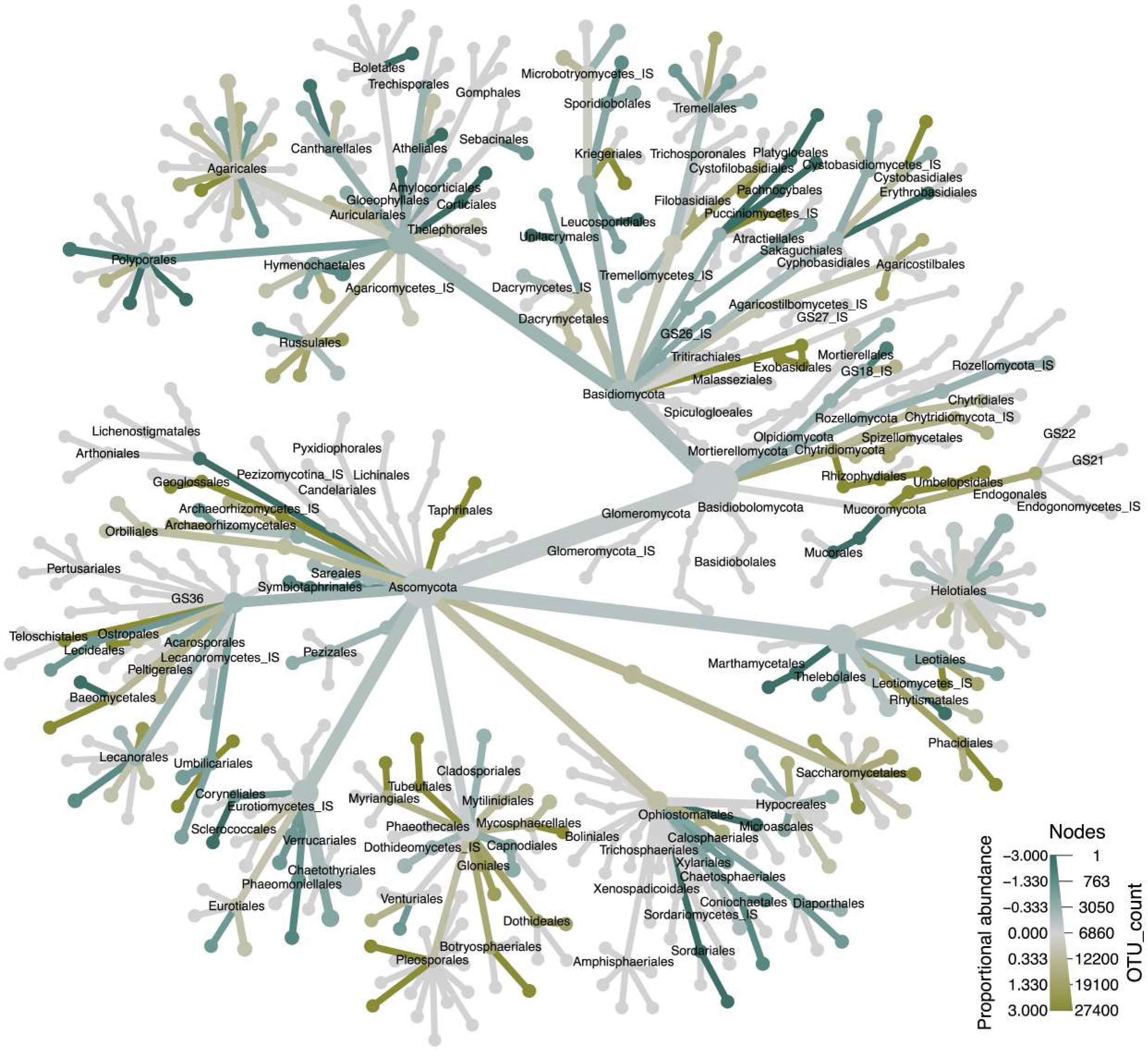
Taxonomic tree of the ITS2 fungal community from all 24 sites, nodes are truncated at Family level for readability and names end at Order level. A tree with names to family level is supplied in supplementary files. Colors represent proportional occurrence rates towards either previously clear-cut (CC) sites in yellow and near-natural (NN) sites in teal.

### Compositional variation among plots

The PERMANOVA analysis revealed a significant long-term effect of clear-cutting on fungal community composition, with an R^2^ of 0.007 (p = 0.001). The NMDS ordination revealed systematic differences in fungal community composition between NN and CC plots with a stress values of 0.167 (Fig. 3a). In nine out of twelve sites, the CC plot shifted in a consistent direction along the ordination axes from the NN pair (Fig 3a). This systematic compositional shift from NN towards CC was in the direction of lower beta-dispersion (i.e., lower variation in species composition between the 20 logs within the plot), lower values of deadwood functional dispersion and evenness, lower volumes of mid-decay stage logs, lower variation in living tree DBH and higher volumes of living spruce (Fig. 2a, Fig. 2b). The ordination also revealed a clear geographical gradient along the second axis that correlated with changes in mean annual temperature and spring snow cover (Fig. 2b).

**Figure 3.**
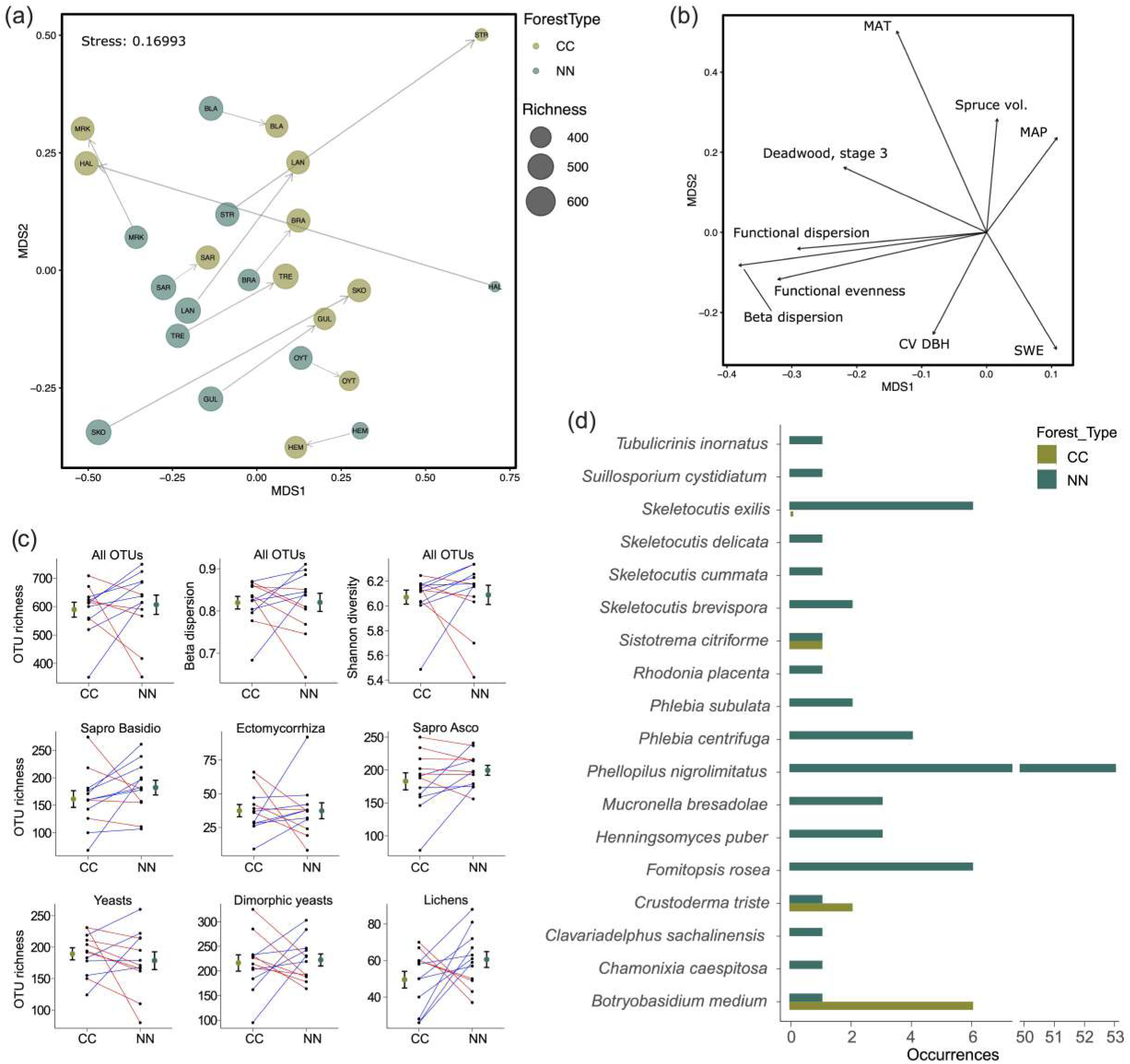
(a) GNMDS ordination of the 24 sites with point size is scaled by OTU richness in each site. Vectors connect the two pairs within each site and point from the NN plot towards the CC plot. (b) Environmental vectors fitted to the ordination in (a). Vectors point in the direction of increase in value and arrow length is scaled by R2 values from the envfit function. (c) Differences between NN and CC sites in richness, beta-dispersion and Shannon diversity, as well as in the OTU richness of the six functional groups of fungi displayed in the OTU ordination in (b). Each line connects to a paired site. Red line means a higher value in CC and blue line means a higher value in NN. (d) Occurrences of all red-listed species and distribution in CC or NN forest. Note the break in x-axis for the very high occurrence of Phellopilus nigrolimitatus.

No significant overall differences in total richness, Shannon diversity, or Beta dispersion were found between CC and NN; instead there was high variation between pairs, with instances of both CC or NN having the higher values among different sites for all indices (Fig. 3c). Similarly, none of the functional groups differed significantly overall in OTU richness (Fig. 3c). However, there was more lichen OTUs in NN plots, but with only weak statistical support (p = 0.06). Dimorphic yeasts were the most species-rich group, followed by yeasts and saprotrophic Ascomycota. The least OTU rich groups were ectomycorrhizal fungi and lichens (Fig 3c).

We identified 18 red-listed species with high bootstrap scores (≥ 0.8) for the assigned species hypothesis (Fig 3d, Fig S5). *Phellopilus nigrolimitatus* was the most common of the red-listed species, found exclusively in NN plots and detected in 53 different logs (Fig 3d). Three of the red-listed OTUs occurred in both CC and NN, while the remaining 15 occurred exclusively in NN. Two of the red-listed OTUs, *Botryobasidium medium* and *Crustoderma triste*, had more occurrences in CC than NN (Fig 3d). Red-listed species were detected in 10 out of 12 NN plots, with the exception of Halden and Hemberget. Hence, the pattern of higher occurrences of threatened species in NN was not driven by just a few plots (Fig S5).

### Dissimilarity models and diversity indices

Functional dispersion of deadwood (i.e. heterogeneity in volumes and characteristics of logs and their decay stages) and yearly mean temperature in the plots, were the only significant predictors related to pairwise dissimilarity in species composition between plots (p < 0.05; Fig. 4a). Of these, only functional dispersion of deadwood was significantly different between CC and NN pairs, with generally higher values in the NN plots, although some NN plots also had among the lowest values (Fig. 1c). Yearly mean temperature increased systematically from north to south in our sampling area reflected in the second axis of the ordination. Functional dispersion of deadwood was significantly associated with both higher OTU richness (p < 0.001, R2 = 0.66, Fig. 4a) and within-transect beta-dispersion ((p < 0.001, R2 = 0.53, Fig. 4a), while yearly mean temperature was not associated with an increase or decrease of any of these community properties (Fig. 4a). The effect of functional dispersion of deadwood was strongest when comparing plots with low dispersion with plots that had medium-to-high dispersion values. Yearly mean temperature, which was also important in structuring community dissimilarity was not significantly correlated with either richness or plot-diversity (Fig. 4a). Beta-diversity, calculated as the Bray-Curtis index between paired plots, was mainly driven by species turnover, not nestedness (Fig. 4b). The mean Bray-Curtis dissimilarity turnover between pairs was 0.53 (range 0.40–0.77), of which the nestedness component made up 0.04 (range 0.01–0.10) (Fig. 4b).

**Figure 4.**
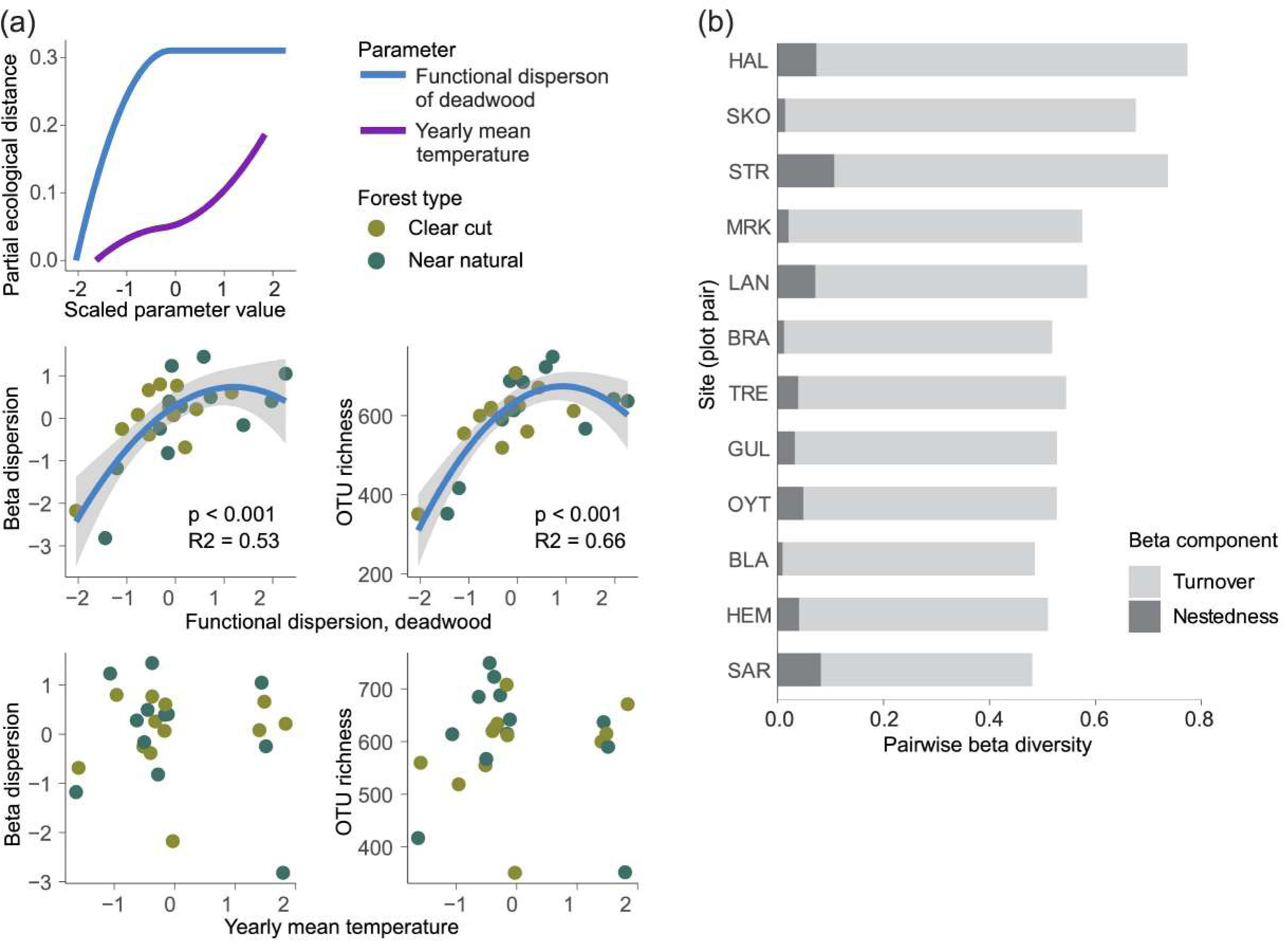
(a) I-splines from a generalized dissimilarity model of the two significant variables predicting the Bray-Curtis distance between any two pairs of sites (top). Scatter-plots of the relationship between beta-dispersion values and OTU richness for each plot and the functional dispersion in deadwood, with associated line of best fit from a linear model (middle). Scatter plots of the relationship between beta-dispersion values and OTU richness for each plot and yearly mean temperature, no significant relationship (bottom). X-axis values are scaled to zero mean and unit variance (b) Beta-diversity partitioned into nestedness and turnover components for each site-pair, calculated as a pairwise Bray-Curtis dissimilarity index.

### Zeta-diversity models

To assess whether rare, intermediate, and common OTUs responded differently to the predictors, zeta diversity tests were conducted. The different orders showed very similar responses to functional dispersion of deadwood and varied in the magnitude of response to the other predictors (Fig. 5). Functional dispersion in deadwood was the most influential predictor (Fig. 5b). The main difference between rare and common orders was the response to DBH variation and snow cover, both were more strongly influencing the common orders (Fig. 5d and e).

**Figure 5.**
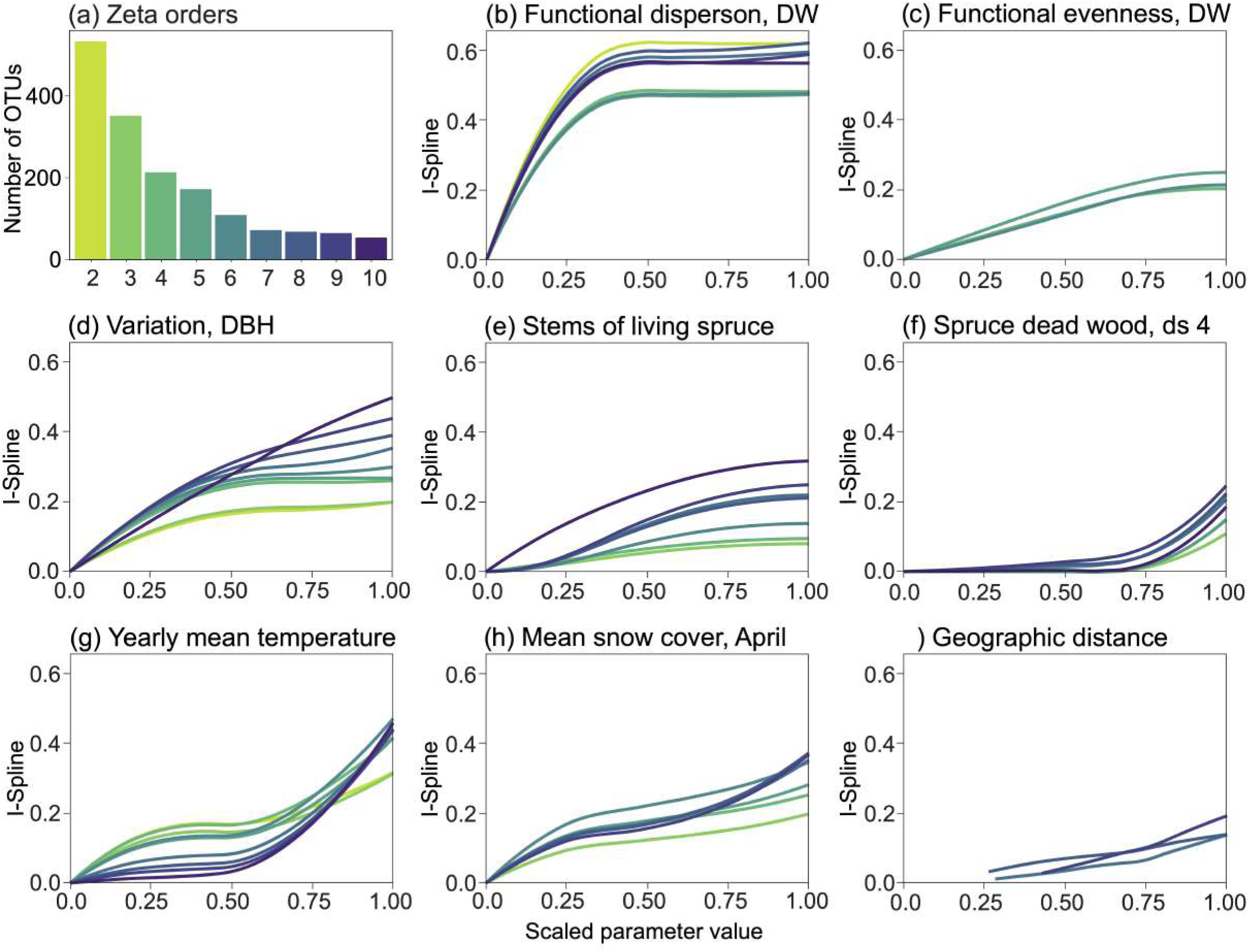
(a) Bargraph of the OTU richness of each zeta-order from 2 to 10. The zeta-orders are the mean number of OTUs shared among any 2 to 10 sites. (b-h) I-splines showing the relative effects of environmental and community variables and on community turnover at the different zeta-orders. I-splines represent the median response after fitting 50 multisite-generalized dissimilarity models with 1000 random selections of site combinations each.

## Discussion

In this study, our main aim was to assess whether there are any long-term consequences of clear-cut forestry on deadwood fungi in boreal spruce forests. We found a significant long term effect of clear-cutting on the overall composition of fungal communities in deadwood. We also found that the near-natural forests tended to be more diverse, but no overall significant difference in OTU richness was found between the forest types. Hence, our first hypothesis – that the near-natural forests would host richer and more diverse communities – was only partially supported. We did, however, observe more red-listed species in the NN plots. This is a well-known pattern in Fennoscandian forests, as many red-listed species are associated with structural components in the forest that are rare or absent in managed stands, for instance, large and old deadwood in the case of *Phellopilus nigrolimitatus,* the most frequent red-listed species we registered and exclusive to NN plots (Stokland and Kauserud, 2004; Tikkanen et al., 2006).

We found functional dispersion of the sampled deadwood to be the main driver of both community dissimilarity between plots, and species richness and diversity within plots. This index describes the variability in deadwood, with high values in plots with variable deadwood (e.g., the simultaneous presence of early, mid, and late decay stages, and small and large volume logs with different characteristics) and low values in plots where the deadwood logs are very similar (Asplund et al., 2025). In essence, we demonstrated that community richness and diversity at plot-scale are more strongly correlated with variation in deadwood and not simply the volume and number of logs at a plot. This result aligns with previous studies based on sporocarp inventory studies of deadwood-associated fungi in boreal forests of Fennoscandia, where it has been found that both amount and variability in deadwood drives the diversity of fungi forming visible sporocarps (Asplund et al., 2026; Juutilainen et al., 2014; Moor et al., 2021; Nordén et al., 2013). Here, we demonstrate that the same pattern holds when considering the whole fungal community through metabarcoding. The patterns were consistent across rare, intermediate and common species (as defined by the zeta-orders), rejecting our fourth hypothesis (H4), namely that rare species would be more strongly affected by changes in forest properties following regeneration after clear-cutting. A previous study in Finland, applying a similar deadwood diversity index, found that richness and abundance of common species depended on a high number of logs at the site and that different decay stages were represented, while the rare (and red-listed) species had higher chance of being present at sites with the presence of large, high volume logs (Hottola et al., 2009). In our study plots, a major difference in properties between CC and NN plots was the lack of large-volume logs in most previously clear-cut forests (Asplund et al., 2024), which at least some rare red-listed Polyporales and Hymenochaetales typically grow on, like *Phellopilus nigrolimitatus* (Stokland and Kauserud, 2004).

Although the overall richness between plots did not differ, there were differences in proportional distribution among several taxonomic groups. We found that, in general, the basidiomycete orders Polyporales and Hymenochaetales were proportionally more abundant in the NN plots. These orders contain many species that are typically recorded in sporocarp surveys and have been found in many studies to be impacted by forestry (Nordén et al., 2013; Stokland and Kauserud, 2004). Helotiales, a highly diverse order in Ascomycota, was also proportionally more abundant in the NN plots. There is likely a high diversity of wood decomposing species in this order that is often overlooked in sporocarp surveys (e.g., van der Wal et al., 2015). In a recent DNA-based study from Finland, it was found that Helotiales species have a higher affinity to naturally fallen dead wood than to artificially made (i.e., felled and placed out into the forest) deadwood (Saine et al., 2024).

Another striking observation was the high diversity of yeasts found in the deadwood, both in Ascomycota (e.g., Saccharomycetales) and Basidiomycota (e.g., Kriegeriales), which mostly had a strong affinity to the CC plots. In contrast, several large groups of dimorphic yeasts (e.g., Chaetothyriales, Tremellales) were more associated with the NN plots. The potential role of yeasts in wood decomposition is not well known, but they have been detected across different decay stages and may contribute to the decomposition process of deadwood (Buzzini et al., 2017; Lara et al., 2014). The yeasts in CC plots may be associated with the large amount of very fresh top-breaks (i.e., tops of trees breaking off during winter snow loads) and may reflect endophytic diversity of yeasts in the living tree (Müller and Hallaksela, 2000; Roll-Hansen and Roll-Hansen, 1980). Noteworthy, one of the prevalent yeast genera we observed, *Scheffersomyces*, is reported to ferment lignocellulose-related sugars within deadwood (Buzzini et al., 2017), further indicating a potential role in decomposition.

We detected a surprisingly high number of lichens inside the deadwood, most of them being proportionally more abundant in the NN plots. Our results mirror results recently reported from Swedish boreal forests (Dahlberg et al., 2025); Similar to their results, we found the genus *Elixia* to be abundantly present inside deadwood alongside species of the genera *Fulgidea, Rhizoplaca*, *Xylospora* and *Hypogymnia* as the most frequently occurring lichenised fungi within our deadwood samples. Notably, during sampling, we carefully shaved off the outer layer of both bark and the underlying wood surface. Thus, it could be speculated that these lichenized fungi could extend their hyphae into the wood and possibly express some saprobic ability.

Despite not finding any differences in richness between CC with NN, we observed that the community composition tended to be different. Hence, we obtained support for our third hypothesis, that the community dissimilarity between CC and NN plots was due to species turnover, and all pairs had very low nestedness values. While many of the conspicuous Polyporales and Hymenochaetales described from numerous other studies discussed above had a clear affinity towards NN plots, saprotrophic basidiomycete fungi only made up a fraction of the species at any of the sites. The common OTUs (higher zeta-orders) was more strongly influenced by the changes in canopy structure, measured as DBH variability and number of stems in each plot, possibly associated with shading, moisture and insolation to the forest floor. These responses are likely indirect, and as they are strongly correlated to the forest type they indicate a long-term shift in the common species that is also driven by the clear-cutting (Stokland and Larsson, 2011). The influence of plot-scale deadwood diversity was the same for both rare and common OTUs and had a similar threshold value for all zeta-orders. This corresponds well to a recent meta-study of deadwood manipulation in managed forests (Sandström et al., 2019). In this meta-analysis, it was found that variation in deadwood was more important than the amount, and similar to what we found, this effect was apparent on the whole community of deadwood fungi, with no specific effect on the rare species (Sandström et al., 2019). It should be noted that the majority of the studies included in the meta-study investigated short-term effects following clear-cutting.

In essence, we found a high beta-diversity between the previously clear-cut and the near-natural forest sites, and this is not dependent on the alpha-diversity (i.e., species richness). Hence, compositional differences among sites (i.e., species turnover) and richness do not necessarily have to be influenced by the same processes (Bratli et al., 2006). We found that variation in deadwood properties within the plot had a high influence on variation in fungal community composition. In addition, increased variation in deadwood was also strongly and positively correlated to the within-plot diversity and the species richness as shown in the generalized dissimilarity models and linear models. This relationship between diversity and richness of deadwood-associated fungi and deadwood variation has been shown in many other studies (Juutilainen et al., 2014; Purhonen et al., 2021; Yang et al., 2021). However, unlike many studies based on sporocarp surveys we did not find general differences in species richness between managed and unmanaged sites. This difference might be related to the DNA-based approach, as more than half of the OTUs in our data are yeasts or dimorphic yeasts and filamentous ascomycete decomposers, which are largely undetected in sporocarp surveys. Moreover, it is possible that our study design may have influenced the richness patterns. By selecting 20 logs per plot, regardless of the total number of logs, we likely under-sampled sites with large amounts of deadwood. This potential bias was partly alleviated by sampling and comparing across several pairs where both the NN and CC have high variation in deadwood amounts, as well as by scaling each plot by the total richness in the dissimilarity models and by including total deadwood volumes per plot as covariates in all the models. While the plot-level species accumulation curves do not systematically saturate at only 20 samples for any plot, the overall CC and NN species accumulation curves overlapped with a slightly higher estimated species richness in NN plots (supplementary data). We can speculate that with equal sampling coverage in all plots, unlike our sampling covering 80-100% of the logs in some sites compared to 18% of the logs in the site with the largest amount of deadwood, we might have discovered a higher total richness associated with increasing amounts of deadwood.

## Conclusion

Our study shows that the fungal community is influenced by clear-cutting several decades later, but that species respond in different ways. Both the composition of rare and common species were affected by the diversity of deadwood. Red-listed species, however, were mostly restricted to the near-natural plots, with only three out of eighteen species occurring in the previously clear-cut plots and more than 90% of the total red-listed occurrences in natural forest, with only three red-listed species also occurring infrequently in CC plots. Our findings highlight that increased habitat availability, even just moderate increases in deadwood volume and heterogeneity, can have a strong effect on community regeneration following clear-cutting. We also show that many understudied groups, like yeasts (e.g., Saccharomycetales) and saprotrophic Ascomycota species (e.g., Helotiales, Sordariales, Dothideales) are diverse and also respond to forestry. The continued practice of clear-cutting and gradual loss of the remaining natural forest is a major threat to the species dependent on the natural forest characteristics. Hence, conservation and forest management practices that promote structural diversity of deadwood, including log size and generating a variety of decay stages, are key for maintaining both rare and common fungal species in boreal forests.

## Supporting information

Supplementary_methods_deadwood_indices

Supplementary_table_deadwood_samples

## Acknowledgements

We acknowledge the Research Council of Norway for financial support to the EcoForest project (No. 320722) and forest owners and municipalities for allowing research on their properties. We acknowledge all EcoForest participants for scientific discussions, especially O. Janne Kjønaas for her contribution to study design and establishing plots. BioFokus and all field assistants are acknowledged for contributions in field work. Michelle Vera Castellanos is thanked for lab assistance in the C/N analyses. We also want to acknowledge Marie David for the illustration used in Figure 1. The bioinformatic analyses were performed on resources provided by Sigma2 - the National Infrastructure for High-Performance Computing and Data Storage in Norway (project NN9338K).

## Data availability

Raw sequences are available at NCBI Short Read Archive (SRA) under accession. OTU tables and predictor variables are available at dataverse.no under the https://doi.org/10.18710/7ZZ4M3.

---

**Figure S1:**
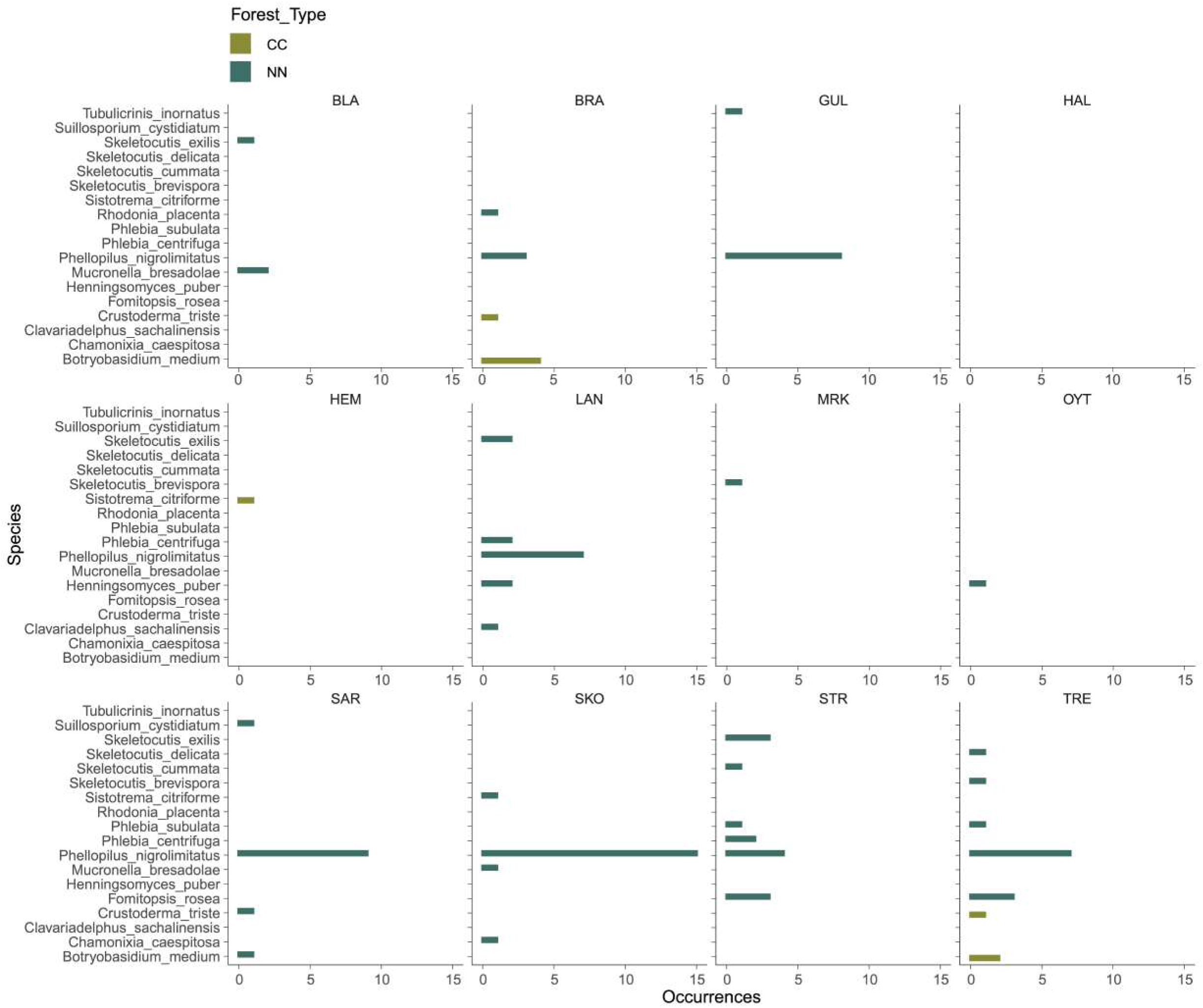
(a) Distribution of volume of sampled logs per decay stage compared to the total volume of logs in each plot. (b) Total number of sampled logs in CC and NN plots per decay stage. (c) log-transformed values of (b).

**Figure S2:**
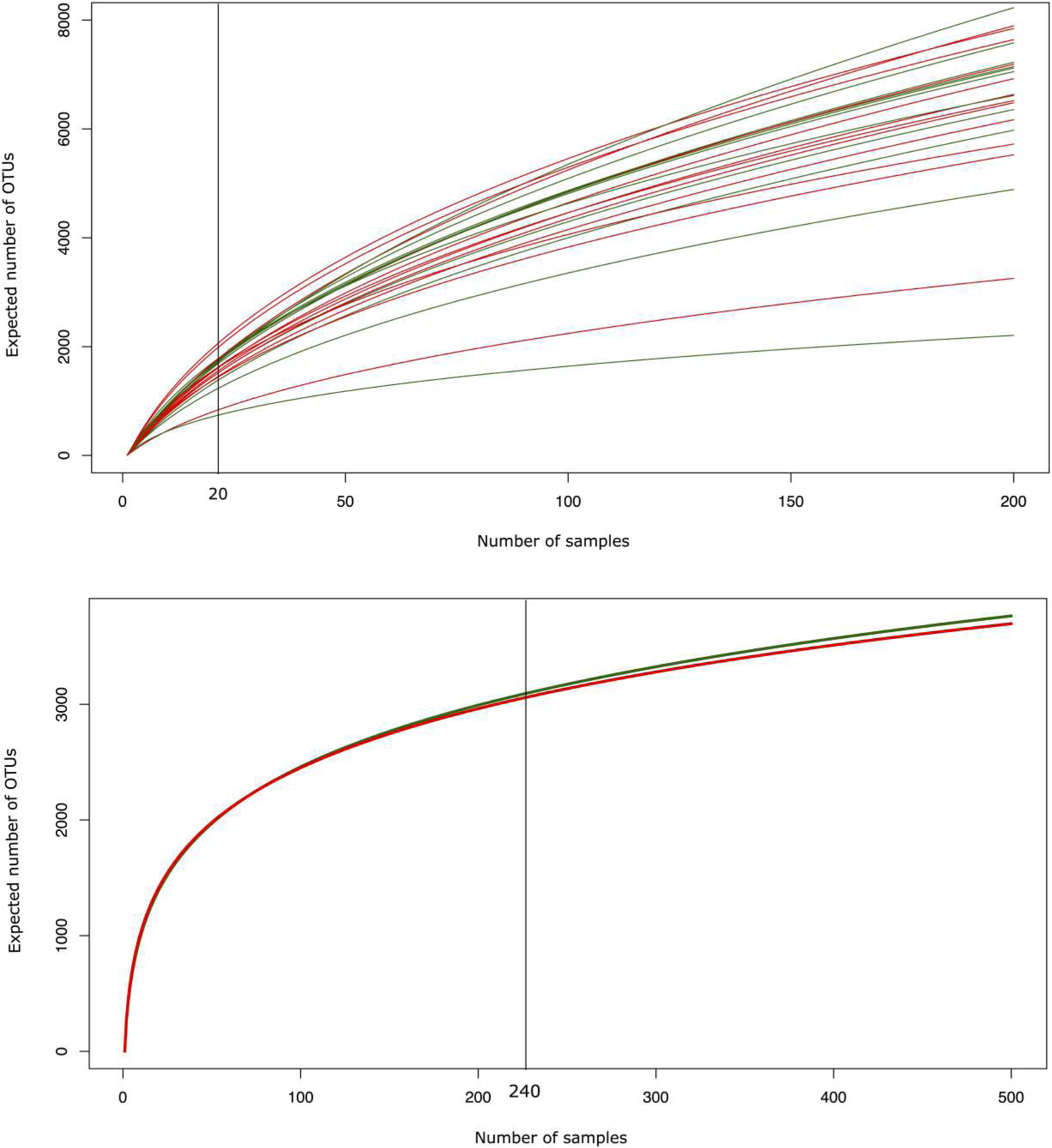
Estimated species accumulation curves for each plot (top) and total for CC and NN samples (below). Horizontal lines at 20 (top) and 240 (below) represent the number of logs we sampled for each plot or in total for the CC and NN.

**Figure S3:**
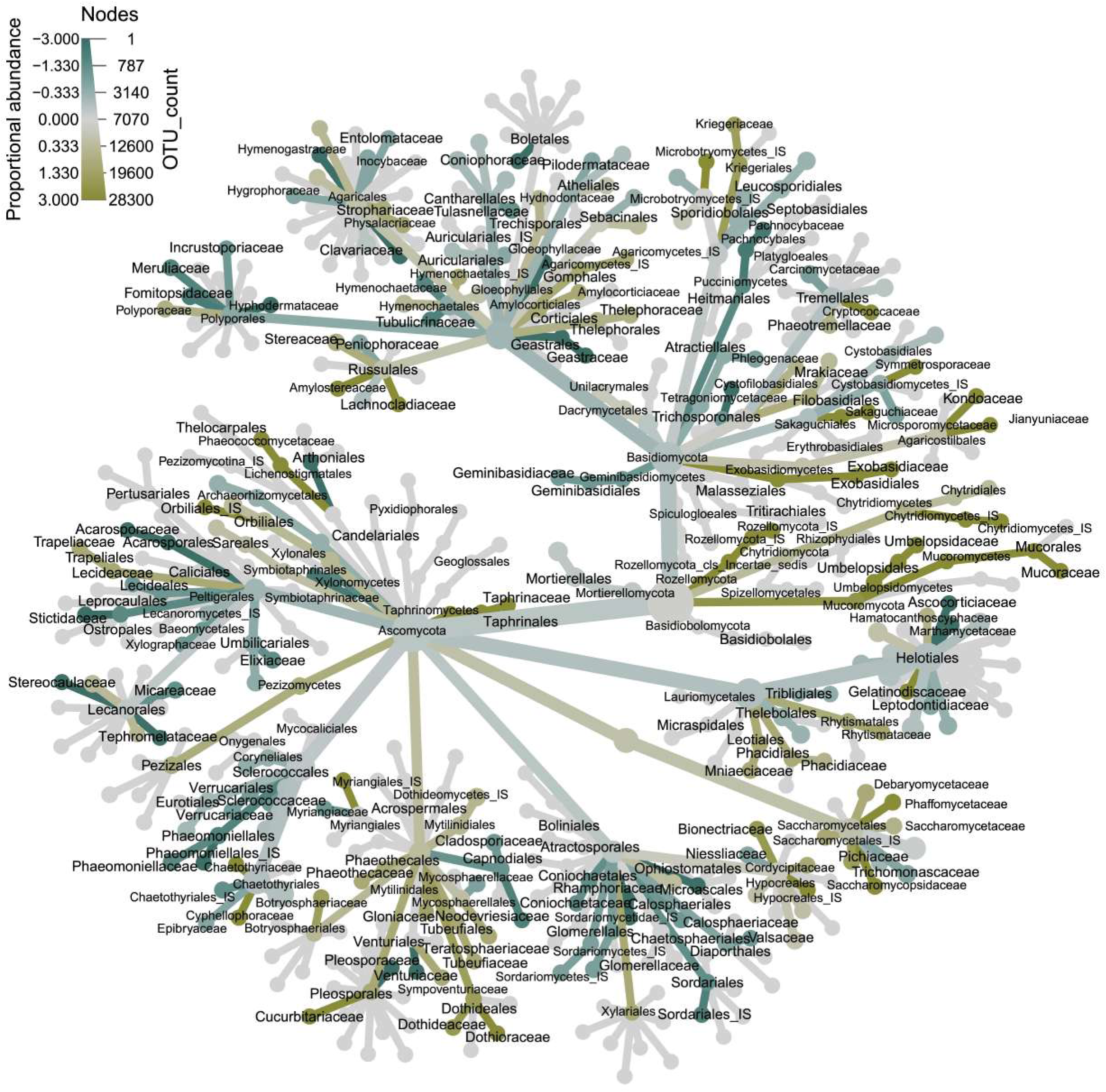
Taxonomic tree of the ITS2 fungal community from all 24 sites, nodes are truncated at Family level. Colors represent proportional occurrence rates towards either previously clear-cut (CC) sites in yellow and near-natural (NN) sites in teal.

**Fig S4:**
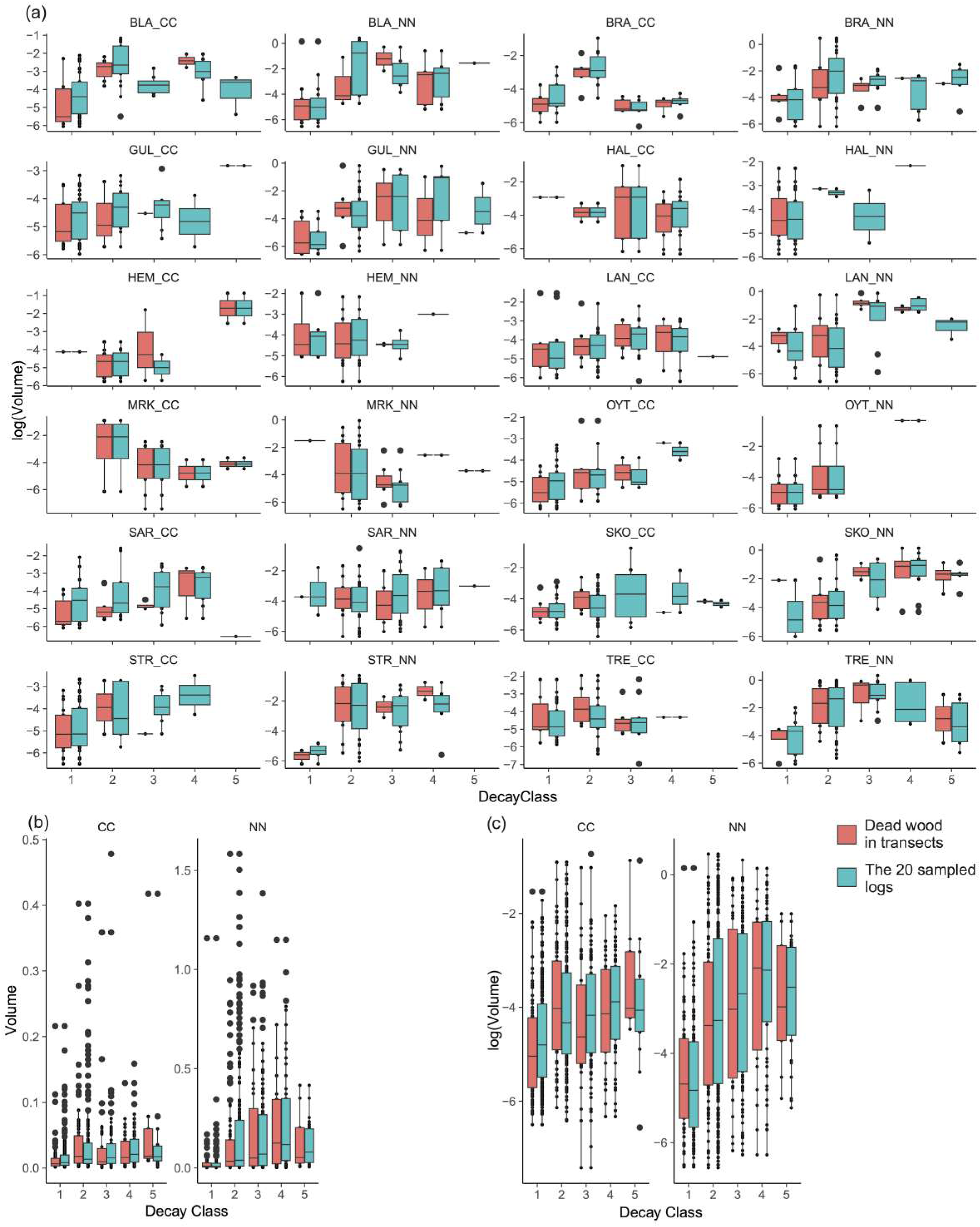
Overview of the occurrences of any of the 18 red-listed species-hypothesis we observed in our data according to which forest site they were found. Color of the bars indicate whether it was found in the NN or CC plot at the site.

