## Supplementary_methods_deadwood_indices for "Long-term shift in community composition of deadwood fungi after clear-cutting"

Functional indices for deadwood

Standard functional diversity indices (FRic, FEve, FDiv, and FDis, Fig 1) were calculated using the R package “mFD” (Magneville et al., 2022). In these calculations, we treated each downed dead wood as if it was a “species” in the calculations and used the continuous variables volume (m3), bark cover (%), epiphyte cover (%), ground contact (%) and the categorical variables, tree species, decay stage class (Viken, 2021), and log type (uprooted, broken or man- made log) as its “traits”. The distribution of deadwood logs with different characteristics is used to represent “species” with different “traits”. We used the *gawdis* package (de Bello et al., 2021) to calculate the optimal weight for each trait when computing the multi-trait dissimilarity measure. Bark cover and epiphyte cover were grouped when weighing the traits, as their high correlation results in a comparable effect on functional diversity calculation, even though they each have different ecological impacts. In the current study, the functional indices used were calculated only on the selection of logs used for eDNA analysis, not the full range of deadwood in the transect. Methods adapted from Asplund et al., (2025)

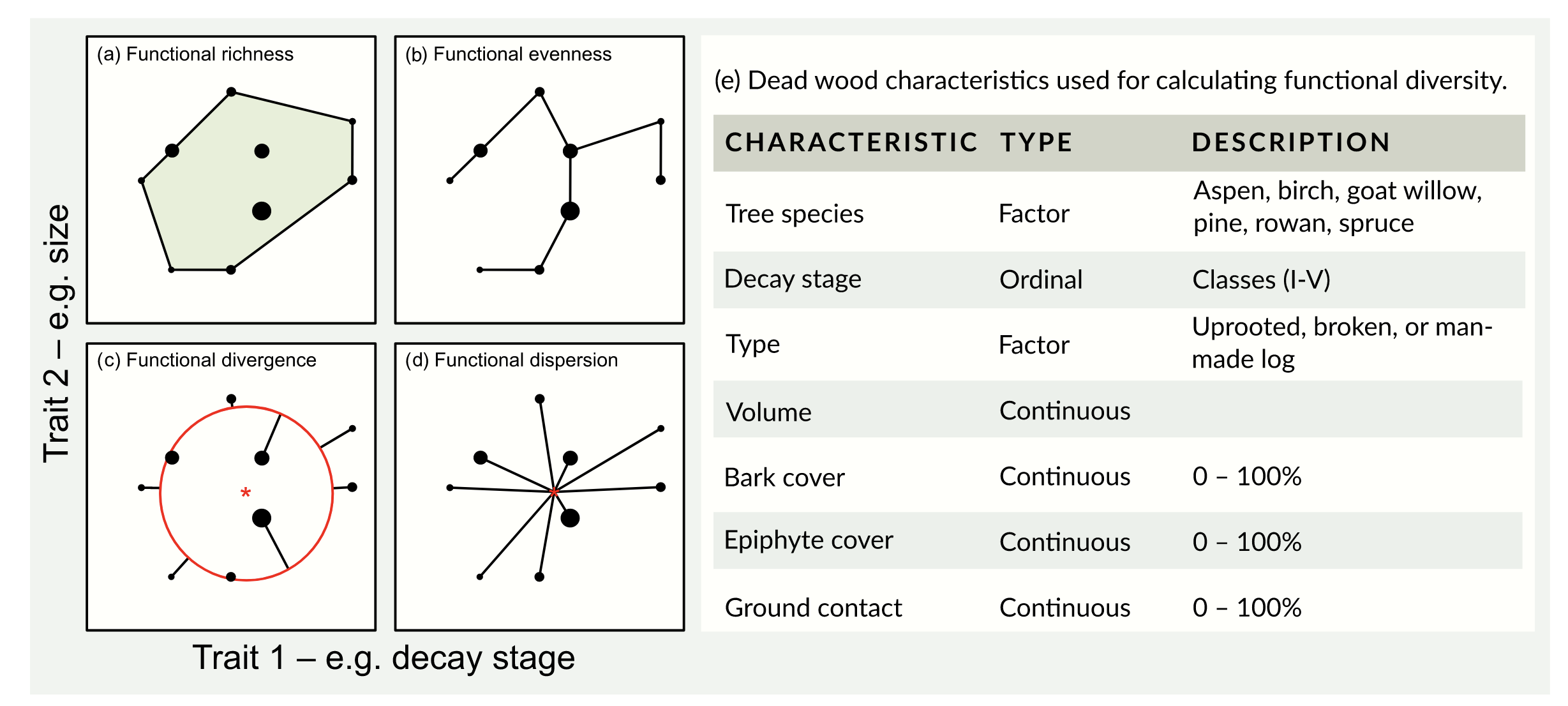

Figure S1: Visual example of the four functional indices. Dots represent a single deadwood unit with values of the traits for each log on the axis. This is a simplified illustration using two traits. In our multivariate analysis, a PCoA was fitted using all the traits explained above (e). (a) Functional richness (FRic) is quantified as the volume of the minimum convex hull that encompasses all the dead wood units within the multidimensional space. (b) Functional evenness (FEve) is measured as regularly distributed the dead wood units are along a minimum spanning tree connecting the units. As such it is a quantification of how regularly the multidimensional space is filled. Functional evenness is highest if the distance between each dot in the minimum spanning tree is equal. (c) Functional divergence (FDiv) is a measure of how much each dead wood unit on average deviates from the average distance to the centre of gravity (red ring), i.e. the average length of the black lines. (d) Functional dispersion (FDis) is calculated as the average distance (black lines) of each dead wood unit to the centre of gravity (red asterisk), where each position is weighted to the number of downed dead wood units. Adapted from Villéger et al., (2008) and Laliberté and Legendre, (2010).

*References*

Laliberté, E., Legendre, P., 2010. A distance‐based framework for measuring functional diversity from multiple traits. Ecology 91, 299–305. https://doi.org/10.1890/08-2244.1

Viken, K.O., 2021. Landsskogtakseringens feltinstruks – 2021. NIBIO.
