## Supplementary_table_deadwood_samples for "Long-term shift in community composition of deadwood fungi after clear-cutting"

*Table S1: Overview of the number of sampled logs in each site and the number of logs that were not sampled.*

| Plot_ID | Sampled | Not Sampled | Sampled proportion |
| --- | --- | --- | --- |
| BLA_CC | 20 | 94 | 17,5438596 |
| BLA_NN | 20 | 20 | 50 |
| BRA_CC | 20 | 34 | 37,037037 |
| BRA_NN | 20 | 59 | 25,3164557 |
| GUL_CC | 20 | 71 | 21,978022 |
| GUL_NN | 20 | 73 | 21,5053763 |
| HAL_CC | 20 | 7 | 74,0740741 |
| HAL_NN | 20 | 13 | 60,6060606 |
| HEM_CC | 18 | 2 | 90 |
| HEM_NN | 20 | 24 | 45,4545455 |
| LAN_CC | 20 | 58 | 25,6410256 |
| LAN_NN | 20 | 40 | 33,3333333 |
| MRK_CC | 20 | 0 | 100 |
| MRK_NN | 20 | 29 | 40,8163265 |
| OYT_CC | 20 | 31 | 39,2156863 |
| OYT_NN | 20 | 11 | 64,516129 |
| SAR_CC | 20 | 50 | 28,5714286 |
| SAR_NN | 20 | 58 | 25,6410256 |
| SKO_CC | 20 | 66 | 23,255814 |
| SKO_NN | 20 | 35 | 36,3636364 |
| STR_CC | 20 | 73 | 21,5053763 |
| STR_NN | 20 | 44 | 31,25 |
| TRE_CC | 20 | 53 | 27,3972603 |
| TRE_NN | 20 | 86 | 18,8679245 |
